# Comparison of algorithms for the detection of enteroviruses in stool specimens from children diagnosed with Acute Flaccid Paralysis

**DOI:** 10.1101/179721

**Authors:** J.A. Adeniji, F. A. Ayeni, A. Ibrahim, K.A. Tijani, T.O.C. Faleye, M.O. Adewumi

## Abstract

With poliovirus eradication within reach, the WHO has included in its recommendations a cell-culture independent algorithm for enterovirus surveillance. This study was designed to compare both the cell culture dependent and independent algorithms and assess how either might impact our perception of the diversity of enterovirus types present in a sample.

Sixteen paired samples (16 isolates from RD cell culture and their corresponding stool suspension. i.e. 32 samples) from AFP cases in Nigeria were analyzed in this study. One of these 16 sample pairs (the control) was previously identified and confirmed as poliovirus 2 (PV-2). All the samples were subjected to RNA extraction, cDNA synthesis, RT-snPCR (the WHO recommended cell-culture independent algorithm) and its modifications for co-infection detection and resolution. Amplicons were sequenced and strains identified using the enterovirus genotyping tool and phylogenetic analysis.

The enterovirus diversity was shown to be the same between RD cell culture isolates and fecal suspension for the control and five (7, 10, 11, 12 & 14) of the samples analyzed. It was however, different for the remaining 10 (62.5%) samples analyzed. Fourteen different enterovirus types were identified in this study. To be precise, 9 (CV-B4, E6, E7, E13, E14, E19, E29, EV-B75 and EV-B77) and 5 (CV-A1, CV-A11, CV-A13, EV-C99 and PV2) EV-B and EV-C types, respectively where detected in this study. It is crucial to mention that E19 and EV-B75were only recovered from RD cell culture isolates while E14, EV-B77, CV-A11 and CV-A13 were only recovered from fecal suspension.

The results of this study show that both the cell culture dependent and independent protocols recommended by the WHO for enterovirus detection unavoidably bias our perception of the diversity of enterovirus types present in a sample. Hence, rather than jettison one for the other, effort should be directed at harmonizing both for increased sensitivity.

## INTRODUCTION

Enteroviruses (EVs) belong to genus *Enterovirus* in the family *Picornaviridae* order *Picornavirales*. There are 13 species in the genus, and the type species of the genus is Species C which has Poliovirus as its best studied member [1]. EVs are non-enveloped viruses with icosahedral capsid symmetry and a diameter of 28-30nM. The genome is an ∼7.5kb, single-stranded polyadenylated, positive-strand RNA with a covalently linked viral protein (VPg) at the 5’ terminus. The single open reading frame (ORF) in the genome is flanked by two untranslated regions (the 5’UTR and 3’UTR). The large polyprotein translated from the single ORF is processed to yield four structural proteins (VP1, VP2, VP3 & VP4) and seven non-structural proteins. The sequence of the VP1 region has been correlated with EV serotype [2] and is now used for identification EV types.

Majority of enterovirus data that has been made available in the last three decades has been courtesy the global polio eradication initiative (GPEI) and specifically their laboratory arm (Global Polio Laboratory Network [GPLN]). Hence, most of these EV isolates (polioviruses [PVs] and nonpolio enteroviruses [NPEVs]) were recovered following the WHO recommended cell culture based enterovirus detection algorithm [3, 4]. With the goal (poliovirus eradication) of GPEI within reach, there is justifiable concern about facility associated escape of polioviruses into the community, post eradication [5]. Hence, as part of the endgame strategy, effort is ongoing to restrict poliovirus research in cell culture globally to few facilities (referred to as essential facilities) with the infrastructure to prevent and contain facility associated escape of the virus [5].

To facilitate implemention of this restriction in the near future, there has been significant motivation to develop very sensitive cell-culture independent strategies for poliovirus (and other NPEVs) surveillance [6 – 8]. In line with this, a cell-culture independent algorithm developed by Nix et al., [6] has been included in the recommended assays for enterovirus detection and identification by the WHO [9]. We recently showed [10] that this WHO recommended cell-culture independent enterovirus detection algorithm [9] misses out enterovirus co-infection. This facilitates underestimation of a very common condition that was instrumental to the circulating vaccine derived poliovirus 2 (cVDPV2) outbreak [11] in Nigeria that lasted almost a decade. Consequently, we have described modification of the assay to expand its capacity, thereby facilitating detection and resolution of coinfection [10, 12].

In the light of the biases [13, 14] and limitations [10, 12] of both the cell culture dependent [4] and independent [9] algorithms, this study was designed to assess the impact of a switch from the former to the latter in the future. Further, it investigated how these algorithms alongside the co-infection (species) resolution assay impact our perception of the diversity of enterovirus types present in a sample. This study finds that both the cell culture dependent [4] and independent [9] algorithms have their strengths and weaknesses and unavoidably bias our perception of the diversity of enterovirus types present in a sample. It demonstrates the need to maximize the benefits of all available strategies in a bid to better describe the diversity of enteroviruses in any sample of interest. Finally, this study documents the first description of a Nigerian strain of EV-B77.

## METHODOLOGY

### Sample Collection

Sixteen RD positive isolates and their corresponding suspensions (making 32 samples in all i.e. 16 pairs of isolate from cell culture and stool suspension) were analyzed in this study. The samples were collected from the WHO National polio laboratory in the Department of Virology, College of Medicine, University of Ibadan, Nigeria (subsequently referred to as the polio lab). Ten of the samples came from five cases (i.e. two stool samples collected at least 24 hours apart from the same case.). The remaining six samples were from six independent cases. One of these six samples was previously identified and confirmed by the Polio Lab as poliovirus 2 (PV-2). All the samples analyzed in this study were collected as part of the National Acute Flaccid Paralysis (AFP) surveillance programme. The samples were collected from children ≤ 15 years presenting with AFP between July and August 2015. The algorithm followed in this study is depicted in Figure 1.

**Figure 1:**
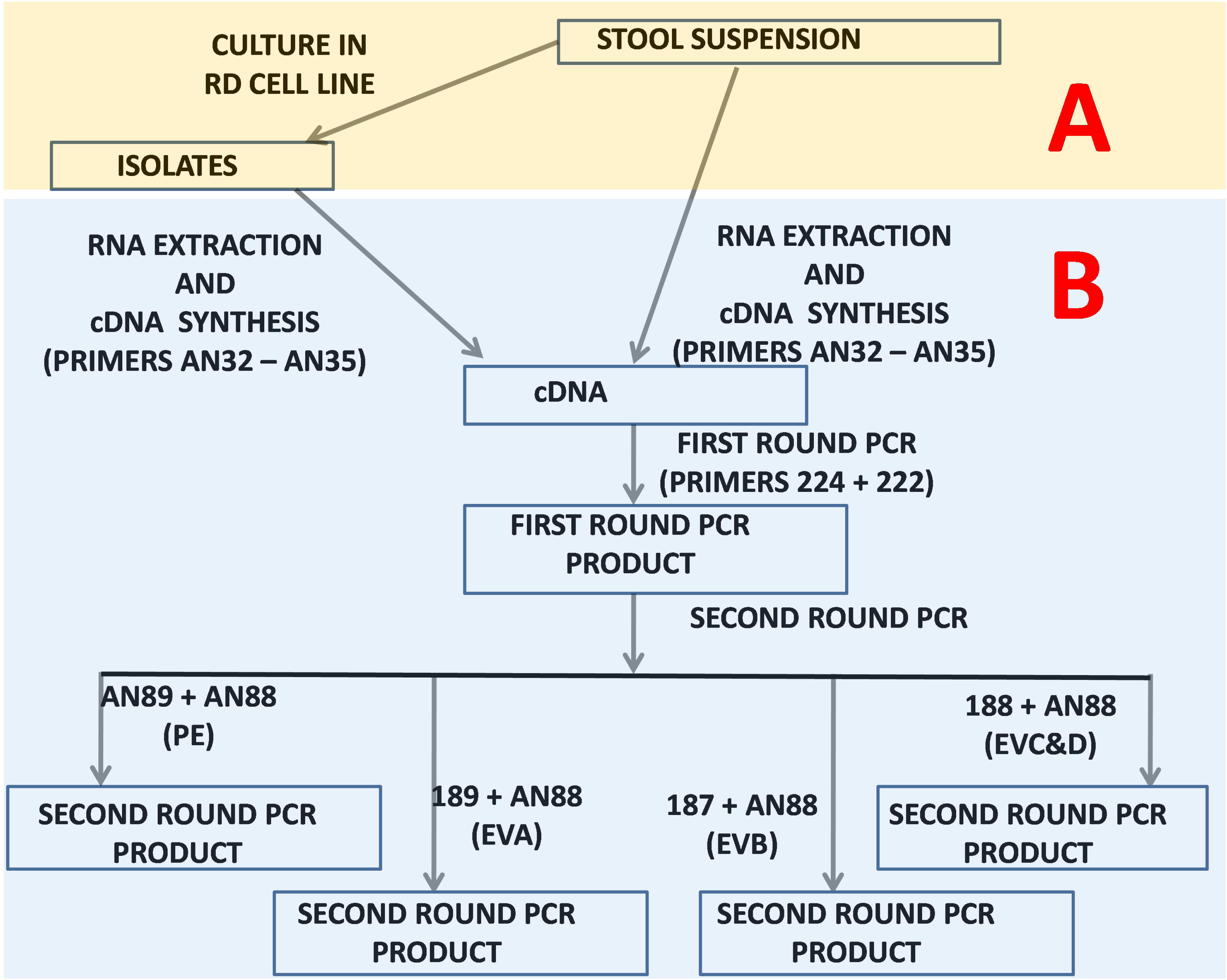
Schematic representation of the algorithm used in this study. (A) Sixteen RD cell culture isolates and their corresponding sixteen fecal suspension were collected from the WHO National Polio Laboratory in Ibadan, Nigeria.. (B) RNA was extracted from all thirty-two (32) samples (RD positive isolates and their corresponding suspension) and subsequently converted to cDNA. The cDNA was used as template in the 1^st^ round PCR assay. The first round PCR assay product was used as template in four different second round PCR assay. Positive samples for the 2^nd^ round PCR assays were sequenced and the result was used for enterovirus identification.

**Figure 2:**
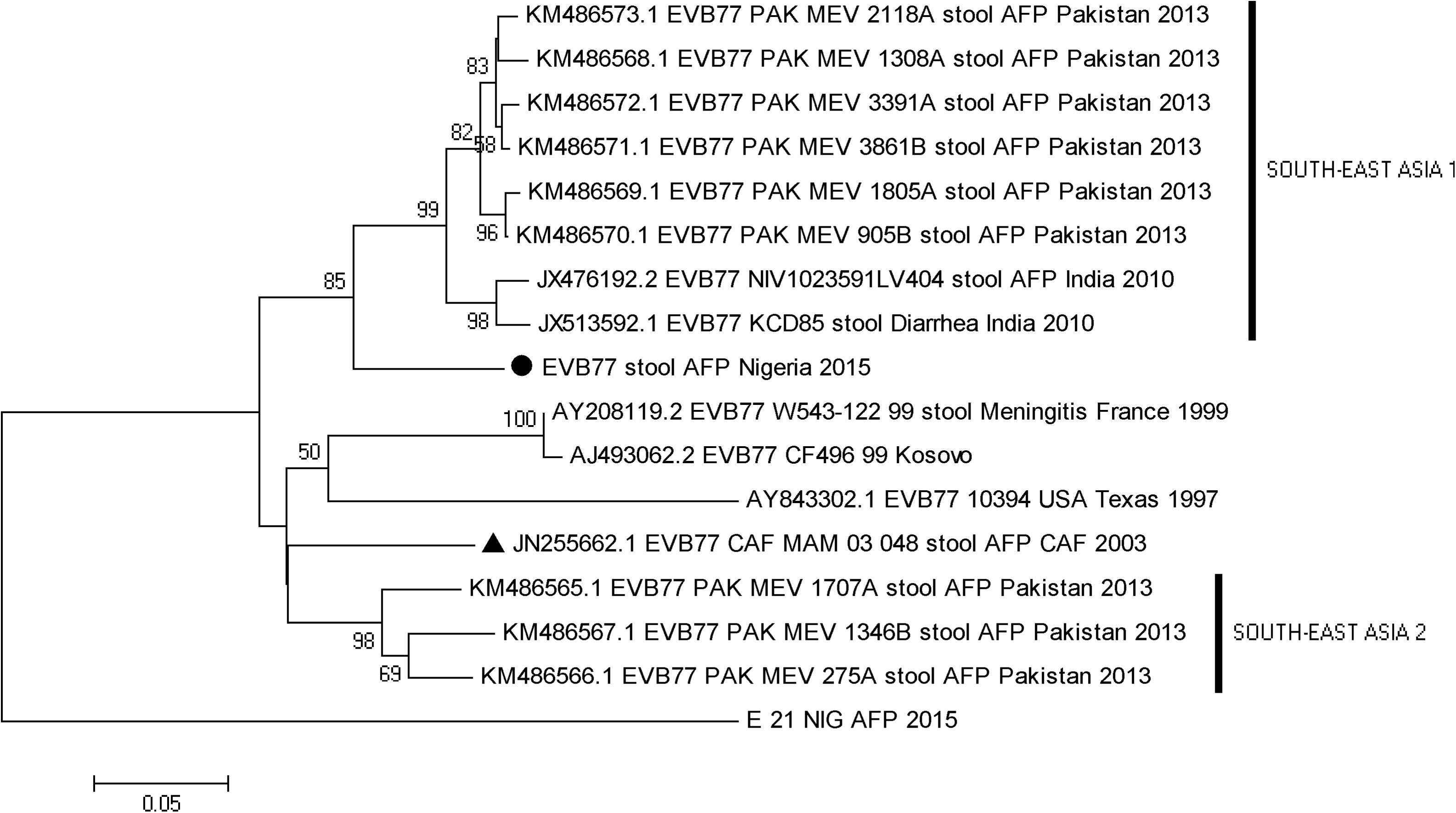
Phylogram of EV-B77. The phylogram is based on an alignment of partial VP1 sequences. The newly sequenced strains are highlighted with Black circle. The strain previously recovered from sub-Saharan Africa in 2003 is indicated with black triangle. The GenBank accession numbers of the strains are indicated in the phylogram. Bootstrap values are indicated if > 50%.

### RNA extraction and cDNA synthesis

RNA was extracted from isolates and suspensions independently using Jena Bioscience Total RNA extraction kit (Jena Bioscience, Jena, Germany) following the manufacturer’s instructions. For cDNA synthesis, Jena Bioscience SCRIPT cDNA synthesis Kit (Jena Bioscience, Jena, Germany) was used according to manufacturer’s instructions. From the extract, 5.25 μL of viral RNA was added to 4.75μL of cDNA synthesis mix. The 4.75μL of cDNA synthesis mix contained 2μL of Script RT buffer, 0.5μL of dNTP mix, 0.5μL DTT stock solution, 0.5μL of RNase inhibitor, 0.25μL of SCRIPT reverse transcriptase and 0.25μL each primers AN32-AN35. The mixture was incubated at 42 °C for 10min followed by 50 °C for 60 minutes in a Veriti thermal cycler (Applied Biosystems, California, USA).

### Polymerase Chain Reaction

The 1^st^ round PCR reaction (Figure 1) was a total of 30 μL reaction. The reaction mix contained 6μL of Red Load Taq, 13.4μL of RNase free water, 0.3μL of primers 224 and 222, and 10μL of cDNA. Thermal cycling was done in a Veriti thermal cycler (Applied Biosystems, California, USA) as follows; 94^°^C for 3 minutes, then 45 cycles of 94^°^C for 30 seconds, 42^°^C for 30 seconds, and 60^°^C for 60 seconds, with ramp of 40% from 42^°^C to 60^°^C. This was then followed by 72^°^C for 7 minutes, and held at 4^°^C until the reaction was terminated.

Four (PE-VP1-PCR, EA-VP1-PCR, EB-VP1-PCR and EC-VP1-PCR) different second round PCR assays were run in this study (Figure 1). The 2^nd^ round PCR assay was also a 30μL reaction. The PCR reaction mix contained 6μL of Red Load Taq, 18.4μL of RNase free water, 0.3μL of forward and reverse primers and 5μL of the first round PCR product. Thermal cycling was done in a Veriti thermal cycler (Applied Biosystems, California, USA). The cycling conditions were 94^°^C for 3 minutes followed by 45 cycles of 94^°^C for 30 seconds, 42^°^C for 30 seconds, and extension at 60^°^C for 30 seconds, with ramp of 40% from 42^°^C to 60^°^C. This was then followed by 72^°^C for 7 minutes and subsequently held at 4^°^C until the reaction was terminated. The PCR products were resolved in a 2% agarose gels stained with ethidium bromide and viewed using a UV transilluminator.

### Amplicon Sequencing

The amplicons of positive PCR reactions for the four second round PCR assays was shipped to Macrogen Inc, Seoul, South Korea, where amplicon purification and sequencing were done. Sequencing was done using the respective forward and reverse primers for each of the four assays. Subsequently, using the enterovirus genotyping tool [15] and the sequence data, the enterovirus genotype and species were determined.

### Nucleotide Sequences Accession Numbers

The sequences obtained from this study have been deposited in GenBank with accession numbers MF686545-MF686568

### Phylogenetic Analysis

The CLUSTAL ***W*** programme in MEGA 5 software [16] was used with default settings to align sequences of the enterovirus type(s) whose Nigerian strains were first described in this study alongside those retrieved from GenBank. Subsequently, a neighbor-joining tree was constructed using the same MEGA5 software [16] with the Kimura-2 parameter model [17] and 1000 bootstrap replicates. The accession numbers of sequences retrieved from GenBank for this analysis are indicated in the sequences name on the phylograms.

## RESULTS

### Polymerase chain reaction (PCR) assay

The expected ∼330bp fragment was successfully amplified for most of the assays carried out. For the *PE-VP1-PCR* screen, of the sixteen RD isolates subjected to this screen, 93.9% (15/16) were positive while 75.0% (12/16) of the corresponding suspensions were also positive. For the *EA-VP1-PCR* screen, 75.0% (12/16) of the RD isolates were positive as were 62.5% (10/16) of the corresponding suspensions. For the *EB-VP1-PCR* screen, 87.5% (14/16) and 68.8% (11/16) of the RD isolates and the corresponding suspensions were positive, respectively. Also, for the *EC-VP1-PCR* screen, 50% (8/16) and 37.5% (6/16) of the RD isolates and the corresponding suspension were positive, respectively (Table 1).

**TABLE 1:** Results of the RT-Semi nested PCR assays done in this study.

| SAMPLE |  | ISOLATES |  |  |  |  | SUSPENSION |  |  |  |  |
| --- | --- | --- | --- | --- | --- | --- | --- | --- | --- | --- | --- |
|  |  | Species Specific Assays |  |  |  |  | Species Specific Assays |  |  |  |  |
| S/N | CASES | PE | EV A | EV B | EV C&D | SUMMARY | PE | EV A | EV B | EV C&D | SUMMARY |
| 1 | Case 1a | + | + | + | + | ++++ | + | + | + | - | +++ |
| 2 | Case 1b | + | - | + | - | ++ | - | - | - | - |  |
| 3 | Case 2a | + | + | + | - | +++ | + | + | + | + | ++++ |
| 4 | Case 2b | + | + | + | + | ++++ | + | + | + | + | ++++ |
| 5 | Case 3a | + | + | + | + | ++++ | + | + | + | - | +++ |
| 6 | Case 3b | + | + | + | + | ++++ | + | + | + | + | ++++ |
| 7 | Case 4a | + | - | + | + | +++ | + | - | + | - | ++ |
| 8 | Case 4b | + | + | + | - | +++ | - | - | - | - |  |
| 9 | Case 5a | + | + | + | - | +++ | - | - | - | - |  |
| 10 | Case 5b | - | - | - | - |  | - | - | - | - |  |
| 11 | Case 6 | + | + | + | - | +++ | + | + | + | - | +++ |
| 12 | Case 7 | + | + | + | + | ++++ | + | + | + | - | +++ |
| 13 | Case 8 | + | - | + | - | ++ | + | - | + | - | ++ |
| 14 | Case 9 | + | + | + | + | ++++ | + | + | + | + | ++++ |
| 15 | Case 10 | + | + | + | + | ++++ | + | + | + | + | ++++ |
| 16 | Control | + | + | - | + | +++ | + | + | - | + | +++ |
| TOTAL SUMMARY |  | 15 | 12 | 14 | 9 |  | 12 | 10 | 11 | 6 |  |

### Enterovirus Genotyping

Of all the sixteen RD isolates, fifteen were amplified, successfully sequenced and typed for the PE-VP1-PCR screen using the enterovirus genotyping tool. Their identities are as follows E7 (3 isolates), E19 (2 isolates), E29 (1 isolate), EV B75 (1 isolate), CV A1 (1 isolate), E6 (2 isolates), E13 (4 isolates). PV2 (1 isolate). For the EA-VP1-PCR screen, twelve RD isolates were successfully amplified but three were successfully typed and their identities are as follows; EV-C99 (1 isolate), CV-A1 (1 isolate), PV2 (1 isolates). For the EB-VP1-PCR screen, fourteen RD isolates were successfully amplified, sequenced and typed and their identities are as follows; E19 (2 isolates), E7 (3 isolates), E6 (2 isolates), E13 (4 isolates), E29 (1 isolate), EV-B75 (1isolate), CV-B4 (1 isolate). For the EC-VP1-PCR screen, nine RD isolates were amplified but two were successfully typed and their identities are; PV2 (1 isolates), EV-C99 (1 isolate) (Table 2). Over all, ten serotypes were identified for the RD isolates PCR screen comprising of specie B (70%) and specie C (30%) (Table 3).

**TABLE 2:** Results of nucleotide sequencing and identification of enterovirus isolates and strains recovered in this study.

| SAMPLE |  | ISOLATES |  |  |  |  | SUSPENSION |  |  |  |  |  |  |
| --- | --- | --- | --- | --- | --- | --- | --- | --- | --- | --- | --- | --- | --- |
|  |  | Species Specific Assays |  |  |  |  | Species Specific Assays |  |  |  |  |  |  |
| S/N | CASES | PE | EV A | EV B | EV C&D | SEROTYPE IDENTIFICATION | PE | EV A | EV B | EV C&D | SEROTYPE IDENTIFICATION | SUMMARY OF SEROTYPES | SPECIES |
| 1** | Case 1a | E19 | NU | E19 | NU | E19 | E13 | NU | E13 |  | E13 | E13, E19 | EV-B* |
| 2 | Case 1b | E19 |  | E19 |  | E19 |  |  |  |  |  | E19 | EV-B |
| 3 | Case 2a | E7 | EV C99 | E7 |  | E7, EV-C99 | NU | EV C99 | NU | EV C99 | EV-C99 | E7, EV-C99 | EV-B, EV-C* |
| 4 | Case 2b | E7 | NU | E7 | EV C99 | E7, EV-C99 | NU | EV C99 | E14 | EV C99 | E14, EV-C99 | E7, E14, EV-C99 | EV-B, EV-C* |
| 5 | Case 3a | E6 | NU | E6 | NU | E6 | E6 | CV A11 | E6 |  | E6, CV-A11 | E6, CV-A11 | EV-B, EV-C* |
| 6 | Case 3b | E6 | NU | E6 | NU | E6 | E6 | CV A13 | E6 | CV A13 | E6, CV-A13 | E6, CV-A13 | EV-B, EV-C* |
| 7 | Case 4a | E13 |  | E13 | NU | E13 | E13 |  | E13 |  | E13 | E13 | EV-B |
| 8 | Case 4b | E13 | NU | E13 |  | E13 |  |  |  |  |  | E13 | EV-B |
| 9** | Case 5a | E13 | NU | E13 |  | E13 |  |  |  |  |  | E13 | EV-B |
| 10 | Case 5b |  |  |  |  |  |  |  |  |  |  |  |  |
| 11 | Case 6 | E7 | NU | E7 |  | E7 | E7 | NU | E7 |  | E7 | E7 | EV-B |
| 12 | Case 7 | E29 | NU | E29 | NU | E29 | E29 | E29 | E29 |  | E29 | E29 | EV-B |
| 13 | Case 8 | EV B75 |  | EV B75 |  | EV-B75 | EV B77 |  | EV B77 |  | EV-B77 | EV-B75, EV-B77 | EV-B* |
| 14 | Case 9 | CV A1 | CV A1 | CV B4 |  | CV-A1, CV-B4 | CV A1 | CV A1 | CV B4 | CV A1 | CV-A1, CV-B4 | CV-A1, CV-B4 | EV-B, EV-C* |
| 15 | Case 10 | E13 | NU | E13 | NU | E13 | E13 | CV A13 | E13 | CV A13 | E13, CV-A13 | E13, CV-A13 | EV-B, EV-C* |
| 16*** | Control | PV 2 | PV 2 |  | PV 2 | PV2 | PV 2 | PV 2 |  | PV 2 | PV2 | PV2 | EV-C |

*NU= NOT USABLE; \*=COINFECTION*

**TABLE 3:** Enterovirus types identified in this study

| ENTEROVIRUS<br>SPECIES | ISOLATE |  | SUSPENSION |  | TOTAL<br>ENTROVIRUS<br>TYPES |
| --- | --- | --- | --- | --- | --- |
|  | ENTROVIRUS<br>TYPES | NUMBER<br>OF<br>TYPES<br>(%) | ENTROVIRUS<br>TYPES | NUMBER<br>OF<br>TYPES<br>(%) |  |
| EV-B | E6, E7, E13,<br>***E19***, E29, ***EV-B75***, CV-B4 | 7 (70%) | E6, E7, E13,<br>***E14***, E29, ***EV-B77***, CV-B4 | 7 (58.3%) | 9 (64.3%) |
| EV-C | EV-C99, CV-A1, PV-2* | 3 (30%) | EV-C99, CV-A1, ***CV-A11***,<br>***CV-A13***, PV-2* | 5 (41.7%) | 5 (35.7%) |
| TOTAL |  | 10 (100%) |  | 12 (100%) | 14 (100%) |

E=Echovirus, EV=Enterovirus, CV=Coxsackievirus, PV=Poliovirus, \*=Control PV2
**Bold and Italicized** = Viruses that were peculiar to the different detection algorithms

Of the corresponding 16 suspensions, twelve (12/16) were amplified but ten (10/16) were successfully sequenced and typed for the PE-VP1-PCR screen and their identities are as follows; E7 (1 strain), E13 (3 strains), E29 (1 strain), EV B77 (1 strain), CV A1 (1 strain), E6 (2 strains), PV2 (1 strain). For the EA-VP1-PCR screen, eight suspensions were amplified, sequenced and typed and their identities are as follows; EV-C99 (2 strains), CV-A11 (1 strain), CV-A13 (2 strains), CV-A1 (1 strain), PV2 (1 strain), E29 (1 strain). For the EB-VP1-PCR screen, twelve amplified but ten were successfully sequenced and typed, and their identities are as follows; E13 (3 strains), E14 (1 strain), E6 (2 strains), E7 (1 strain), CV-B4 (1 strain), E29 (1 strain), EV-B77 (1 strain). For the EC-VP1-PCR screen, six suspensions were successfully amplified, sequenced and typed and their identities are; EV-C99 (2 strains), CV-A13 (2 strains), PV2 (1 strain), CV-A1 (1 strain) (Table 2). Overall, twelve serotypes were identified the suspension PCR screen comprising of specie B (58.3%) and specie C (41.7%) (Table 3).

The enterovirus diversity was shown to be the same in the control (S/N 16) and 33.3% (5/15) of the samples analyzed (Table 2). To be precise, the diversity of enteroviruses was the same between RD cell culture isolates and fecal suspension for the control (S/N 16), cases 4a, 5b, 6, 7 and 9 (Table 2). The diversity of enteroviruses was however, different between RD cell culture isolates and fecal suspension for the remaining 66.7% (10/15) of the sample pairs analyzed (Table 2).

In summary, fourteen different enterovirus types were identified in this study. To be precise, Nine (CV-B4, E6, E7, E13, E14, E19, E29, EV-B75 and EV-B77) and Five (CV-A1, CV-A11, CV-A13, EV-C99 and PV2) EV-B and EV-C types, respectively where detected in this study (Table 3). It is essential to emphasize that the single PV2 detected in this study was the control provided by the Polio Lab.

### Phylogeny of EV-B77

This is the first EV-B77 strain described in Nigeria and the second in sub-Saharan Africa till date. The topology of the phylogenetic tree suggests that the EV-B77 detected in this study is different from all that has been described till date. More importantly, it is different from the only sub-Saharan Africa strain described till date which was recovered in Central Africa Republic in 2003.

## DISCUSSION

### Direct detection from clinical specimen vs after culture in RD cell line

From this study, it was observed that more enteroviruses were detected per sample by the PE-VP1-PCR assay after the suspension had been subjected to culture in RD cell line. For example, Case 1b (Table 2) isolate was identified as E19 while there was no evidence of enterovirus presence in the corresponding suspension. In the same light, the isolates of cases 4b and 5a (Table 2) were identified as E13 while there was also no evidence of enterovirus presence in their corresponding suspensions. Considering, enteroviruses were detected in both the isolate and stool suspensions of other samples, and even the ≥ 24-hour pair of some of the samples in questions, it is unlikely that the observation is due to the presence of nonspecific inhibitors of PCR. Rather, this finding suggests that in the fecal suspension, the virus titre might have been too low (i.e. below the detection limit of the assay) to be detected directly. However, RD cell culture appeared to increase the virus titre to a level that was subsequently detectable by the PE-VP1-PCR assay. This thereby validates the value and use of cell culture for enterovirus detection and identification as it significantly increases virus titre and thereby enhance our capacity to detect and identify the virus types present.

It is however important to note that though some EV types were recovered in both the fecal suspension and RD cell culture, some types appear to be specifically recovered in each detection algorithm (Table 3). The enterovirus diversity was shown to be the same between RD cell culture isolates and fecal suspension for the control and 5/15 (cases 4a, 5b, 6, 7 and 9) of the sample pairs analyzed. It was however, different for the remaining 10/15 (66.7%) sample pairs analyzed. Particularly fascinating is the observation that in some instances the enterovirus isolate detected by the PE-VP1-PCR assay in RD cell culture supernatant is different from what was detected in the corresponding suspension. For example, in case 9 (Table 2), the isolate was identified as EVB75 while EVB77 (first detection in Nigeria) was identified in the corresponding suspension. Also, in case 1a (Table 2) where the isolate was identified as E19, E13 was detected in the corresponding stool suspension, despite the fact that it is well known [18,19] and also documented in this study (Table 2) that RD cell line is both susceptible and permissive to E13. Hence, if the most abundant genome was selectively detected in the above stated instances, these discrepancies suggest that in either case the most abundant genome in the suspension was different from that in the cell culture supernatant. This therefore confirms that culture in RD cell line selectively amplifies one enterovirus genome over the other in cases of co-infection [13, 14, 20], even in cases where both enterovirus types belong to the same species (Table 2). Should this observation be a valid biological phenomenon, its biological basis might further illuminate how culture of enteroviruses in RD cell line influences our perception of the serotype diversity in a sample. Furthermore, this might also indicate that the dynamics of enterovirus culture in RD cell line might not be representative of what happens in the intestinal tract and consequently should not be represented as such.

### The impact of mixture resolving assays

Cases of enterovirus co-infection were established in 53.3% (EV-B/C=40%: EV-B=20%) of the samples analyzed in this study. It is however worthy of note that in these co-infected samples (Table 2), the enterovirus types identified with the PE-VP1-PCR assay were mainly EV-Bs while the EV-C co-infection was majorly detected by the species specific assays. The only exception was in Case 9, were both in the fecal suspension and the RD cell culture isolate, the enterovirus type detected using the PE-VP1-PCR assay was CVA1 while the EVB-VP1-PCR assay detected CVB4 (Table 2). The perceived predilection of the PE-VP1-PCR assay for EV-Bs is not because the primers used for the assay have a bias for EV-Bs. In fact, similar studies using the same assay directly on fecal suspensions without culture in RD cell line show an abundance of EV-As [21]. While those where the same assay was used directly on fecal suspensions that did not show CPE in RD cell line, showed an abundance of EV-Cs [12]. Hence, the predominance of EV-Bs as documented by the PE-VP1-PCR assay, in this study, might be due to the fact that only samples that had yielded isolate in RD cell lines were selected and analyzed in this study. Considering the EV-B bias [13, 14, 20] of RD cell line, this might not be surprising. This observation however suggests that in cases of co-infection involving different enterovirus species, the chances of detecting all the enterovirus present in the sample will be more likely enhanced by the addition of species specific primers to the PCR protocols. For, members of the same species, however, combining cell culture with direct detection from the specimen might be the strategy of choice (Table 2).

### The value of paired samples

The need and value of collecting two stool samples (paired samples) about 24 hours apart from any AFP case is well entrenched in the GPEI enterovirus detection protocols [4, 9]. The results of this study further emphasize the importance of this principle for enterovirus surveillance. For example, it was observed that E13 and E19 were detected in Case 1a but only E19 was identified in Case 1b. Also while E7 & EVC99 were detected in Case 2a, E14 in addition to E7 & EVC99 were detected in Case 2b. More importantly, E13 was detected in Case 5a, while no enterovirus was detected in Case 5b (Table 2). The results of this study therefore further demonstrate that without paired samples, many enterovirus infections would be missed. Consequently, we recommended that this principle be implemented for enterovirus surveillance in general and not just for AFP surveillance.

### Enterovirus detection algorithms and the risk of facility associated escape of poliovirus post containment

We have shown that both the cell culture dependent [4] and independent [9] protocols recommended by the WHO for enterovirus detection unavoidably bias our perception of the diversity of enterovirus types present in a sample (Tables 2 & 3). We have also shown the shortcomings of a PanEnterovirus RT-PCR detection assay which is predicated on the false assumption that co-infections are not significant when enterovirus infections are being considered. Though the anticipated need to prevent the risk of facility associated escape of polioviruses cannot be overemphasized, the findings of this study suggest that enterovirologists should attempt to maximize the benefits of available strategies in a bid to better describe the diversity of enteroviruses in any sample of interest.

On the other hand, effort should be put into expanding the species [22] or serotype [23] specific nextgen sequencing strategies that have already been developed to accommodate other enterovirus types and species. They also have to be expanded to go beyond using isolates recovered from cell culture to direct detection from clinical specimen. Such development might facilitate a successful switch from cell culture dependent to independent strategies without necessarily losing out on breadth and sensitivity.

## AUTHOR CONTRIBUTIONS

1. Study Design (FTOC, AMO, AJA)
2. Sample Collection, Laboratory and Data analysis (All Authors)
3. Wrote, revised, read and approved the final draft of the Manuscript (All Authors)

## ACKNOWLEDGEMENTS

We thank the WHO National Polio Laboratory in Ibadan, Nigeria for providing the anonymous isolates analyzed in this study. This study was done before the global withdrawal of OPV2 in April 2016. This study was funded by contributions from the authors.

